# Prefrontal neural ensembles develop selective code for stimulus associations within minutes of novel experiences

**DOI:** 10.1101/2020.08.26.269035

**Authors:** Kaori Takehara-Nishiuchi, Mark D Morrissey, Maryna Pilkiw

## Abstract

Prevailing theories posit that the hippocampus rapidly learns stimulus conjunctions during novel experiences, whereas the neocortex learns slowly through subsequent, off-line interaction with the hippocampus. Parallel evidence, however, shows that the medial prefrontal cortex (mPFC, a critical node of the neocortical network supporting long-term memory storage) undergoes rapid modifications of gene expression, synaptic structure, and physiology at the time of encoding. These observations, along with impaired learning with disrupted mPFC, suggest that mPFC neurons may exhibit rapid neural plasticity during novel experiences; however, direct empirical evidence is lacking. We extracellularly recorded action potentials of cells in the prelimbic region of the mPFC, while male rats received a sequence of stimulus presentations for the first time in life. Moment-to-moment tracking of neural ensemble firing patterns revealed that the prelimbic network activity exhibited an abrupt transition within a minute after the first encounter of an aversive but not neutral stimulus. This network-level change was driven by ~15% of neurons that immediately elevated their spontaneous firing rates and developed firing responses to a neutral stimulus preceding the aversive stimulus within a few instances of their pairings. When a new sensory stimulus was paired with the same aversive stimulus, about half of these neurons generalized firing responses to the new stimulus association. Thus, prelimbic neurons are capable of rapidly forming ensemble codes for novel stimulus associations within minutes. This circuit property may enable the mPFC to rapidly detect and selectively encode the central content of novel experiences.

## Introduction

Encoding of new information results from changes in neural responses to inputs through activity-dependent modifications of synaptic strength (Hebb, 1949; Kandel and Spencer, 1968). Prevailing theories posit that rapid synaptic plasticity takes place within hippocampal circuits during a novel experience, which guides the subsequent, gradual formation of its memory trace in the neocortex over weeks (McClelland et al., 1995; O’Reilly and Rudy, 2001; Frankland and Bontempi, 2005; Winocur et al., 2010). Consistent with this view, the connectivity of CA3-CA1 synapses in the hippocampus is strengthened upon a single aversive event (Whitlock et al., 2006; Broussard et al., 2016), and neurons in CA1 region develop selective firing patterns for a specific location (*i.e*., place field) within a few minutes of exploration (Bittner et al., 2015, 2017). In contrast, in the medial prefrontal cortex (mPFC), a critical node of the neocortical network supporting long-term memory storage (Frankland and Bontempi, 2005; Tonegawa et al., 2018; Takehara-Nishiuchi, 2020a), neuron ensembles strengthen the selectivity for stimulus associations (Takehara-Nishiuchi and McNaughton, 2008; Kitamura et al., 2017; Morrissey et al., 2017) and object-context association (Weible et al., 2012) over weeks after learning. The weeks-long neural plasticity is accompanied by gradual modifications of local circuits that include NMDA-receptor-dependent synaptic reinforcement (Takehara-Nishiuchi et al., 2006) and changes in synaptic structures (Restivo et al., 2009; Vetere et al., 2011; Kitamura et al., 2017).

The slow, weeks-long circuit modifications in the mPFC, however, do not negate the possibility that the initial step of these changes might start at the time of encoding. In fact, one hour after training in contextual fear conditioning, the transcription of multiple plasticity-related genes is upregulated in the mPFC, leading to immediate changes in synaptic structure and physiology (Bero et al., 2014). Parallel studies also showed that by enhancing the activity of mPFC neurons during training, it is possible to facilitate the acquisition of hippocampus-dependent memories (Benn et al., 2016; Volle et al., 2016; Jarovi et al., 2018; Shibano et al., 2020). These observations, along with earlier findings of impaired hippocampus-dependent memories with disrupted mPFC (Hannesson et al., 2004; Takehara-Nishiuchi et al., 2005; Zhao et al., 2005; Barker and Warburton, 2008; Gilmartin and Helmstetter, 2010; Devito and Eichenbaum, 2011; Gilmartin et al., 2013; Bero et al., 2014), led us to hypothesize that the mPFC may work with, but not follows, the hippocampus to form new memories (Takehara-Nishiuchi, 2020b).

To address this possibility, we investigated how neuron ensembles in the mPFC change firing patterns, while rats underwent trace eyeblink conditioning for the first time in life. In this task, subjects associate a neutral conditioned stimulus (CS) with an aversive unconditioned stimulus applied near the eyelid (US) over a short temporal gap (Woodruf-Pak and Disterhoft, 2008; Takehara-Nishiuchi, 2018). In humans, the formation of CS-US association is contingent on conscious awareness of their temporal relationship, justifying its use as a model of declarative memory (Clark and Squire, 1992). Acquisition of the association depends on the hippocampus (Moyer et al., 1990; Weiss et al., 1999) and the mPFC (Weible et al., 2000; Takehara-Nishiuchi et al., 2005), while with time, its expression becomes dependent on the mPFC (Kim et al., 1995; Takehara et al., 2003). In the first conditioning session, the CS evokes significant firing rate changes in some neurons in the dorsal hippocampus (Berger et al., 1976; Hattori et al., 2015) and anterior cingulate and prelimbic regions of the mPFC (Weible et al., 2003; Hattori et al., 2014). It remains unknown, however, whether these CS-responding neurons represent sensory features of the CS or its higher-order, relational features, such as CS-US associations. This distinction is essential because even in well-trained rats, only ~30% of CS-responding prelimbic neurons differentiate their firing responses depending on whether the CS is predictive of the US or not (Takehara-Nishiuchi and McNaughton, 2008; Morrissey et al., 2017). Thus, the critical question is whether prelimbic neurons start to develop the selectivity for the CS-US association *during* the first session.

## Materials and Methods

### Animals

All experiments were performed on seven male Long-Evans rats (Charles River Laboratories, St. Constant, QC, Canada) weighing 440–600 g at the time of surgery. The data from one rat (Rat 1) was collected as a part of our previous study (Morrissey et al., 2017). Rats were housed individually in Plexiglass cages and maintained on a reversed 12 hr light/dark cycle. Water and food were available *ad libitum*. All methods were approved by the Animal Care and Use Committee at the University of Toronto (AUP 20010962).

### Electrodes for single-unit recording

Tetrodes were made in-house by twisting together four polyimide-coated nichrome wires (Sandvik, Stockholm, Sweden) following our previous work (Takehara-Nishiuchi and McNaughton, 2008; Morrissey et al., 2017). To independently adjust depths, each tetrode was housed inside a screw-operated microdrive. The complete microdrive-array consisted of a bundle of 14 microdrives, each guiding a tetrode, contained within a 3D printed plastic base (Kloosterman et al., 2009). The microdrive-array also enclosed the Electrode Interface Board (EIB-54-Kopf, Neuralynx, Bozeman, MT, United States) to which all electrodes were connected and served as the interface between the recording and stimulating electrodes and the recording system. Prior to implantation, the impedance of the nichrome tetrode wires was reduced to 200-250 kOhms by electroplating electrode tips with gold. Tetrodes were then drawn inside a 7×2 bundle (~2×0.5 mm) made of polyimide tubing. A drop of sterilized mineral oil was added at the microdrive-array base to ensure the smooth movement of the tetrodes after implantation.

### Surgical procedures

The microdrive-array was implanted above the prelimbic region of the medial prefrontal cortex with the same procedure as those used in our previous work (Takehara-Nishiuchi and McNaughton, 2008; Morrissey et al., 2017). All surgeries were conducted under aseptic conditions in a dedicated surgical suite. Rats were anesthetized with isoflurane (induced at 5% and maintained at 2–2.5% by volume in oxygen at a flow rate of 0.9-1.0 L/min; Halocarbon Laboratories, River Edge, NJ, United States) and placed in a stereotaxic holder with the skull surface in the horizontal plane. In Rat 1, a craniotomy was opened at 3.2 mm anterior and 1.4 mm lateral to bregma, and the dura matter removed. The microdrive array was then lowered at a 9.5° medial angle until the base made contact with the surface of the brain. In Rats 2–7, a craniotomy was opened between 3.3–5.4 mm anterior and 0.65–.95 mm lateral to bregma. After removal of the dura mater, the array was placed at the brain surface (1.5-1.75 mm ventral from the skull surface) at a 15° forward angle. Rats 2–7 were also implanted with microinfusion cannulae bilaterally in the lateral entorhinal cortex, which were used to collect data that were not reported in this paper. The craniotomy was covered with Gelfoam (Pharmacia & Upjohn, NJ, United States), and the array was held in place with self-adhesive resin cement (3M, MN, United States) and self-curing dental acrylic (Lang Dental Manufacturing, Wheeling, IL, United States) with screws inserted in the skull. Two additional ground screws were placed above the parietal cortex. During the surgery, all tetrodes were lowered 1.0–1.5 mm into the brain. For the next seven days (Rats 2–7) or four weeks (Rat 1), the rat was connected to the recording system daily to monitor the quality of neural signal and tetrode movement. During this period, each tetrode was gradually lowered (0.03–1 mm per day) to target tetrode tips to the prelimbic region at 2.0– 3.6 mm ventral from the brain surface. Two reference electrodes were positioned superficially in the cortex (1 mm below the brain surface).

### Behavioral paradigm and data acquisition

All rats experienced the same general experimental procedure. During the first two days, rats were placed in the conditioning apparatus without receiving any stimulus presentations. Rat 1 was placed in a large dark rectangular box, fitted with an LED light source and speaker. Within the box, the rat was enclosed in a square Plexiglas container (20×20×25 cm), fitted with holes on one side to enable sound-waves from the speaker to enter the enclosure. The experimental apparatus for Rats 2–7 consisted of a 50×50×50 cm black box, fitted with a sliding door connected to a small hallway and a 6×12 cm perforated Plexiglas window. A speaker and a 5 mm LED were attached outside of the window. On the third day, the conditioning began, and this paper reports neuronal activity recorded on this day. The conditioned stimulus (CS) was presented for 100 ms and consisted of an auditory stimulus (85 dB, 2.5 kHz pure tone) or a visual stimulus (white LED light blinking at 50 Hz). The unconditioned stimulus (US) was a 100 ms mild electrical shock near the eyelid (100 Hz square pulse, 0.3–2.0 mA), and the intensity was carefully monitored via an overhead camera and adjusted to ensure a proper eyeblink/head turn response (Morrissey et al., 2012; Volle et al., 2016). The timing of the CS and US presentation was controlled by a microcomputer (Arduino Mega, Arduino, Italy). The US was generated by a stimulus isolator (ISO-Flex, A.M.P.I., Jerusalem, Israel) and applied through a pair of stainless steel wires chronically implanted in the eyelid. The CS presentations were separated by a random interval ranging from 20 to 40 seconds.

The recording session consisted of two epochs of trace eyeblink conditioning, each of which included 70-100 trials of CS presentations. The two epochs were separated by a 10 min rest period. Each epoch began with 20-37 presentations of the CS by itself, followed by 50-80 trials in which the CS was paired with the US, separated by a stimulus-free interval of 500 ms (Figure 1A). The first epoch included only one of the two CS (*e.g*., auditory CS), and the second epoch included the other CS (*e.g*., visual CS). Before and after each epoch, the rat was placed in a comfortable rest box (Rat 1) or the hallway separating the conditioning chambers (Rats 2–7).

**Figure 1.**
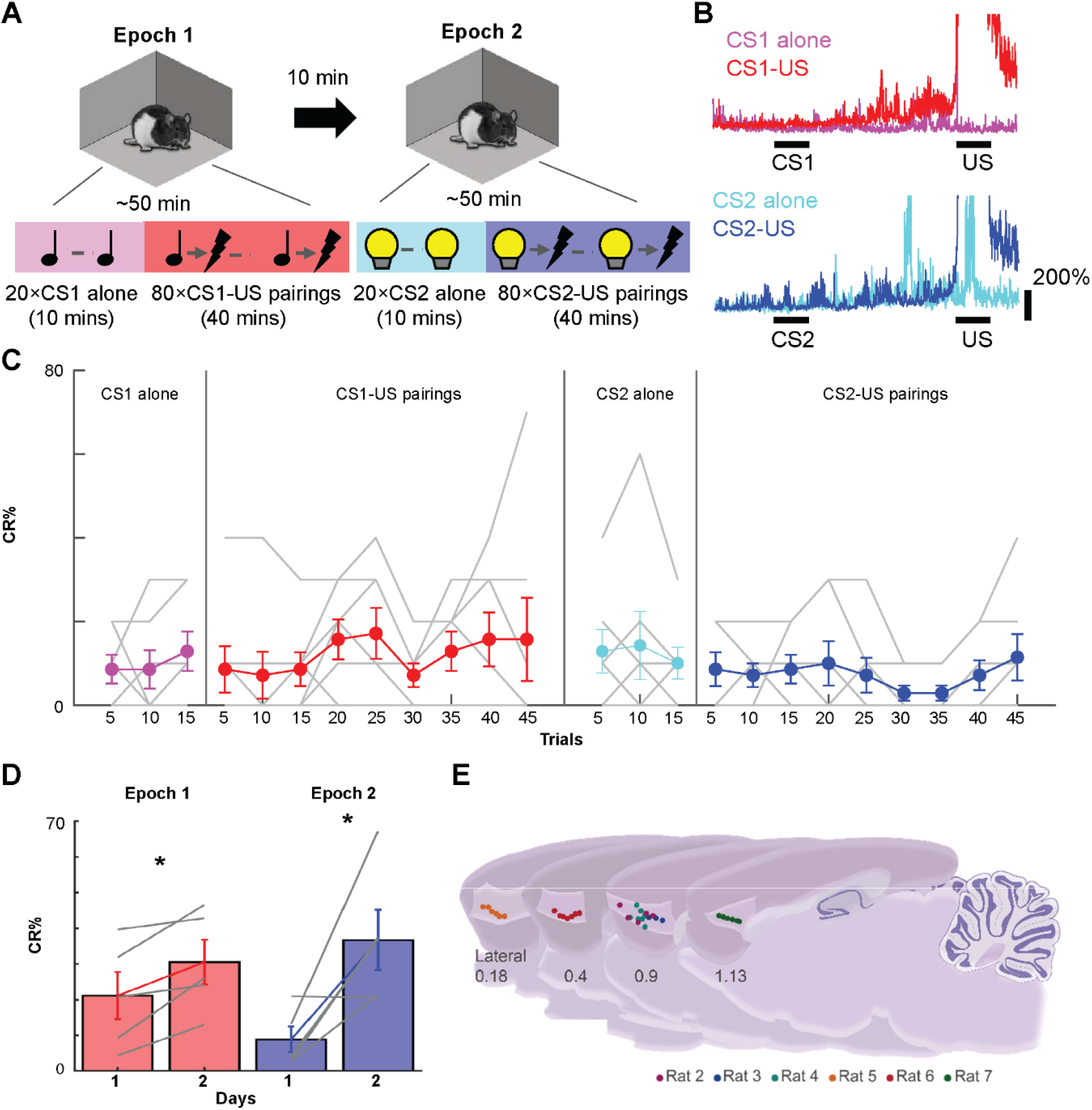
Behavioral paradigm and recording location. ***A***, Seven male rats underwent two epochs of conditional associative learning task in the same chamber. In each epoch, a tone or light was presented alone (CS-alone, 20 trials) or preceding the mildly aversive eyelid shock (US) by 500 ms (CS-US, 50-80 trials). ***B***, Amplitudes of CS-evoked eyelid muscle activity in a representative rat. The muscle activity was normalized by the activity during the period before CS onset. The increased muscle activity toward US onset indicates anticipatory blinking responses (CRs). ***C***, The proportion of trials in which the rats expressed CRs (CR%, mean ± s.e.m.) in a series of 10 trials with an increment of 5 trials. Gray lines indicate the data in each rat. ***D***, CR% in CS-US blocks on the first and second day (mean ± s.e.m.) *, *p* < 0.05, paired t-test. ***E***, Recording locations. Each dot indicates the location of a tetrode tip confirmed in the histological analysis. The brain of Rat 1 was sectioned coronally, which makes it difficult to depict on the same diagram. However, the location of tetrodes in this rat was comparable to the other rats (see Morrissey et al., 2017).

During the conditioning session, we simultaneously recorded action potentials from individual neurons in the right prelimbic cortex and electromyogram (EMG) activity from the left upper eyelid. Action potentials were captured using the tetrode technique, which allows for recording the activity of many individual neurons per recording session (Wilson and McNaughton, 1993). Rats were connected to the recording system through an Electrode Interface Board (EIB-54-Kopf, Neuralynx, Bozeman, MT, United States) contained within the microdrive array fixed to the animal’s head. The EIB was connected to a headstage (Rat 1, HS-54, Neuralynx, Bozeman, MT, United States; Rat 2-7, CerePlex-M 64, Blackrock Microsystems, UT, United States), and signals were acquired through the Cheetah Data Acquisition System (Rat 1; Neuralynx, Bozeman, MT, United States) or the Cerebus Neural Signal Processor (Rat 2-7; Blackrock Microsystems, UT, United States). A threshold voltage was set at 40–75 mV, and if the voltage on any channel of a tetrode exceeded this threshold, activity was collected from all four channels of the tetrode. Spiking activity of single neurons was sampled for 1 ms at 30-32 kHz, and signals were amplified and filtered between 600–6,000 Hz. EMG activity was continuously sampled at 6,108 Hz and filtered between 300–3,000 Hz.

### Behavior analysis

To detect conditioned responses (CR), EMG activity was analyzed with custom codes written in Matlab (Mathworks, Natick, MA, United States). The instantaneous amplitude of the EMG signal was calculated as the absolute value of the Hilbert transform of the signal (using the *hilbert* function in Matlab). In each trial, we calculated EMG amplitude during two 500-ms windows, one from CS offset (CR_value) and the other from 1 sec before CS onset (Pre_value). A trial was removed if its Pre_value was more than three scaled medial absolute deviations away from the median across all trials. A threshold was set as a mean ± four standard deviation of Pre_value. A trial was judged to contain the CR if CR_value was larger than the threshold. The proportion of CS-US trials with the CR in the total valid trials (CR%) was used to quantify the progress of associative learning. It was calculated either in a series of 10 trials with an increment of 5 trials on the first day (Figure 1C) or in all valid trials on the first and second day (Figure 1D). Two rats were removed from the latter calculation due to the poor quality of EMG recording on the second day.

### Neural activity analyses

#### Data preprocessing

Putative single neurons were isolated off-line using a specialized software package in Matlab (KlustaKwik, author: K.D. Harris, Rutgers, The State University of New Jersey, Newark, NJ; MClust, author: D.A. Redish, University of Minnesota, Minneapolis, MN; Waveform Cutter, author: S.L. Cowen, University of Arizona, Tucson, AZ, United States). Both automatic spike-sorting and manual sorting were used to assign each action potential to one of the neurons recorded simultaneously on one tetrode based on the relative amplitudes on the different tetrode channels and various other waveform parameters including peak/valley amplitudes, energy, and waveform principal components. The final result was a collection of timestamps associated with each action potential from a given neuron. Only neurons with <1% of inter-spike intervals distribution falling within a 2-ms refractory period were used in the final analysis.

#### Ensemble transition analysis

Spiking activity of simultaneously recorded neurons during the entire duration of two epochs was used for the analysis. It was binned into a series of 1-ms bins and then smoothed over 20-ms hanning windows. Then, they were binned into 1-s bins. The binned firing rates were converted to Z scores with their mean and standard deviation and stored in a firing rate matrix (Neurons×Time bins, Master matrix). To eliminate the direct impact of trials on the network activity, bins including stimulus presentations were removed from the analysis. To quantify the degree of differentiation of ensemble firing patterns before and after a trial, we calculated Mahalanobis distances between two sets of ensemble firing vectors. This measure allows for quantifying the ensemble selectivity without averaging firing rates at fixed time points and successfully applied to ensemble recording data in the previous studies (Hyman et al., 2012; Rozeske et al., 2018). From the ensemble firing rate matrices, two matrices of 60 columns (covering ~1-min window) were extracted: one matrix is from immediately before the first trial of each CS presentation type (PreMat), and the other after the trial (Post_Mat_). The generalized Mahalanobis distance (DMah) between two matrices before and after the trial (between-block comparison) was defined as:

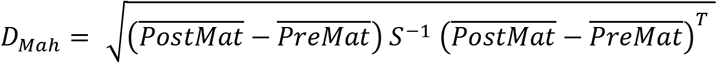

where 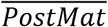 is the mean of the matrix across time bins, and S^-1^ is the inverse of the pooled covariance matrix, defined as

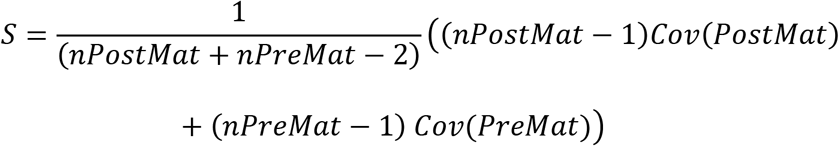

where *Cov* refers to the covariance matrix of the input matrix, and *nMat* refers to the number of time bins included in the input matrix. As a control, we also extracted the third matrix (ContMat) from the time window adjacent to the PreMat (for the CS-US trial) or the PostMat (for the CS-alone trial). D_mah_ was then calculated between the two adjacent matrices (within-block comparison). We were not able to generate PreMat in Rats 2-4 because the first CS1-alone trial was presented less than one minute after the start of the recording. Therefore, these rats were removed from the calculation of D_mah_ for the CS1-alone trial and CS2-alone trial.

#### Neuron categorization

Principal component analysis (PCA) was performed on the Master matrix. Among all principal components (PCs), we selected a PC with the greatest difference in the value between 1-s windows before and after the first CS1-US trial. The PC became high upon the trial in four rats, while it became low in three rats. In the latter case, the sign of the loadings was flipped. Each neuron has a unique loading for the PC. Based on the loadings, neurons were categorized into three groups: ones with the loadings greater than or equal to 0.15 (positive-loadings neurons), ones with the loadings smaller than or equal to 0.15 (negative-loadings neurons), and the remaining neurons (neutral-loadings neurons). These criteria were chosen based on the visual inspection of the distribution of loadings, which was significantly different from the normal distribution (Figure 2A; Lilliefors test, *P* < 0.05).

**Figure 2.**
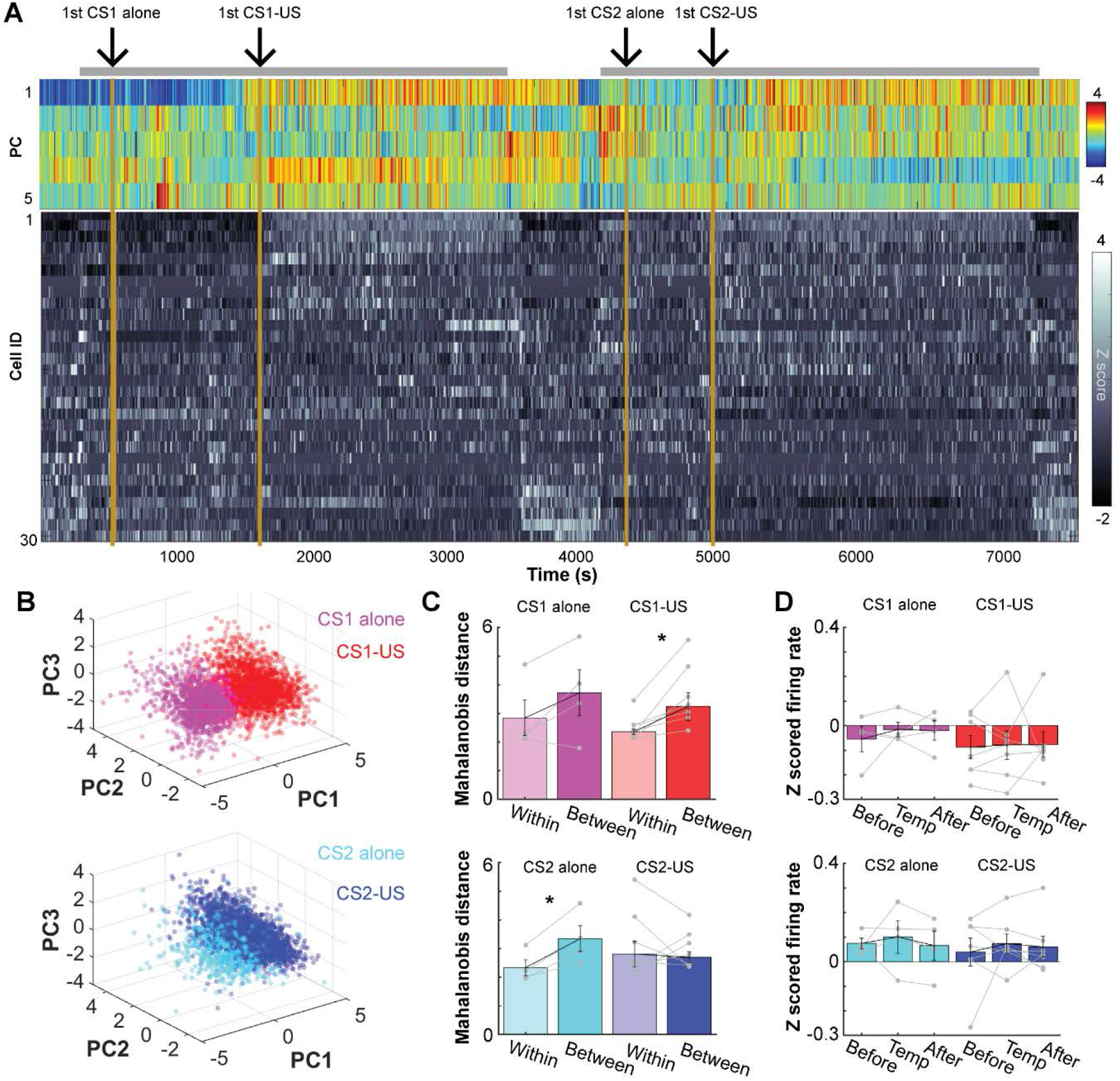
Abrupt transition of the prefrontal network state upon the first encounter of an aversive stimulus. ***A***, Top five principal components (PC; top) of z-scored, 1-sec binned firing patterns of simultaneously recorded neurons (bottom) during the entire recording period including the stimulus presentations in a box (grey bar) and a rest period outside the box. ***B***, 3D projections of PC1-3 in A. ***C***, Mahalanobis distances of the ensemble activity (mean ± s.e.m. and data in individual rats) between two adjacent time windows. They were taken from periods before and after the first trial of each type (Between) or two adjacent periods before or after the first trial (Within). *, *p* < 0.05, Wilcoxon signed-rank test. ***D***, Normalized firing rates averaged across all neurons in each rat (mean ± s.e.m. and data in individual rats) during three windows used to calculate Mahalanobis distances in C.

#### Single-neuron firing rate analysis

To examine stimulus-evoked firing patterns in each trial (Figures 3B, C), firing rates in each neuron were calculated in a series of 100-msec bins and converted into z scores by using the mean and standard deviation of binned firing rates during a 1-s period before CS onset. CS-evoked firing responses were defined as the mean of z-scored firing rates across six 100-ms bins after CS onset. Spontaneous firing rates were defined as raw firing rates during 600-ms windows starting from 1 second before CS onset (Figure 3D).

**Figure 3.**
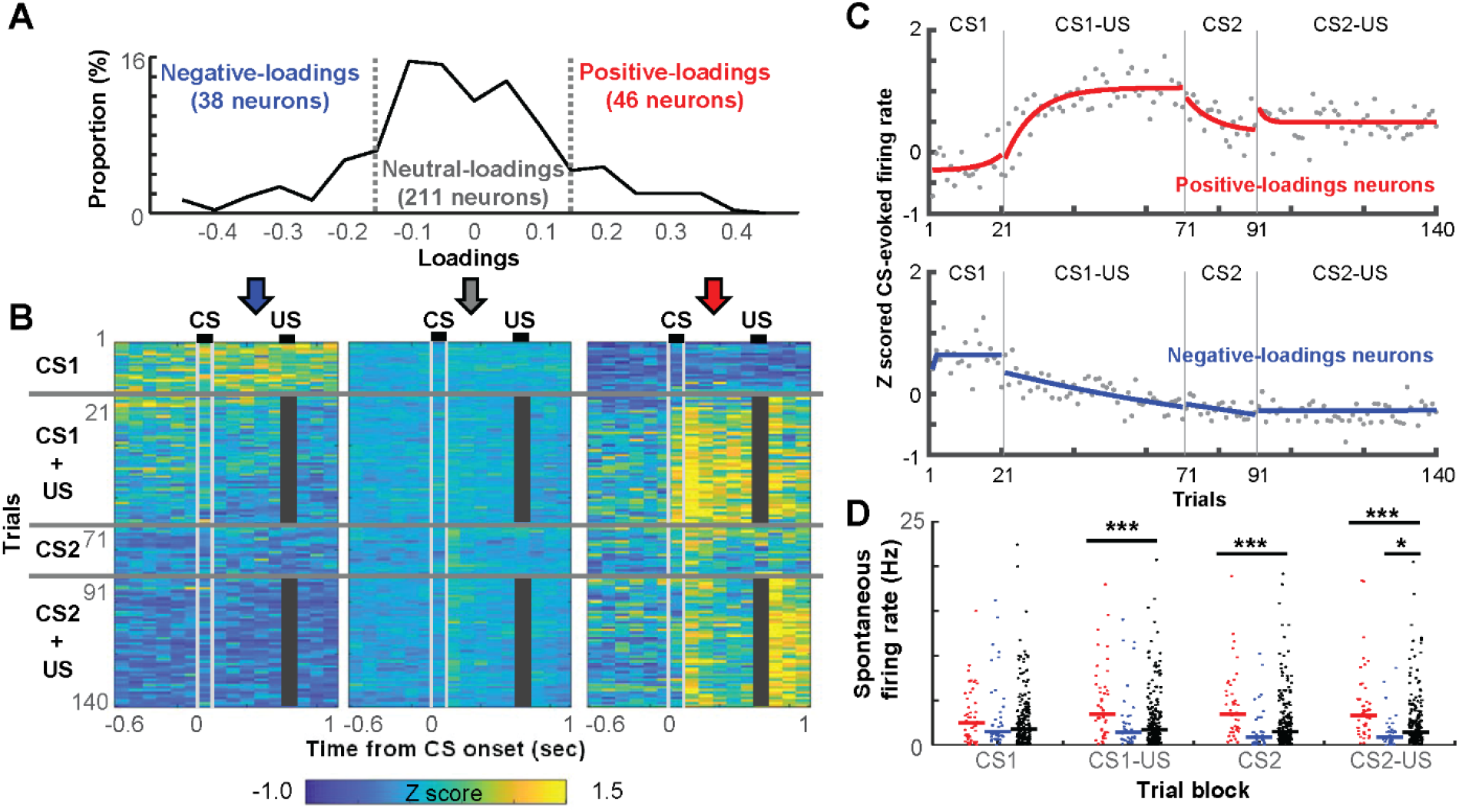
The rapid development of selective ensemble firings for stimulus associations. ***A***, The distribution of loadings on the principal component capturing the network transition after the first aversive stimulus. ***B***, Averaged, 100-ms binned, z-scored firing rates in each trial in neurons positive-loadings (≥ 0.15), negative-loadings (≤ −0.15), as well as the remaining, neutralloadings neurons. Grey bars mask the shock artifact. ***C***, Z-scored, CS-evoked firing rates averaged across all positive- or negative-loadings neurons. Each dot depicts the firing rate in one trial. By fitting an exponential curve (red, blue) to the data, we estimated the number of trials that were required to reach an asymptote. ***D***, Spontaneous firing rates (a dot for each neuron, lines show the median) in the block of four different trial types. *, *p* < 0.05; ***, *p* < 0.001, Kruskal-Wallis test, *post hoc* Dunn’s test.

To quantify the degree to which each neuron changed firing rates after the presentation of the CS or US (Figure 4), firing rates were averaged across trials during the CS (a 600-ms window from CS onset to US onset; FR_CS), US (a 300-ms window starting from 100 ms after US offset; FR_US), and Baseline (a 600-ms window starting from 1 second before CS onset; FR_Pre). Then, the Response Index (RI) was calculated as

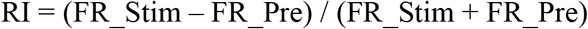

where FR_Stim was either FR_CS or FR_US.

**Figure 4.**
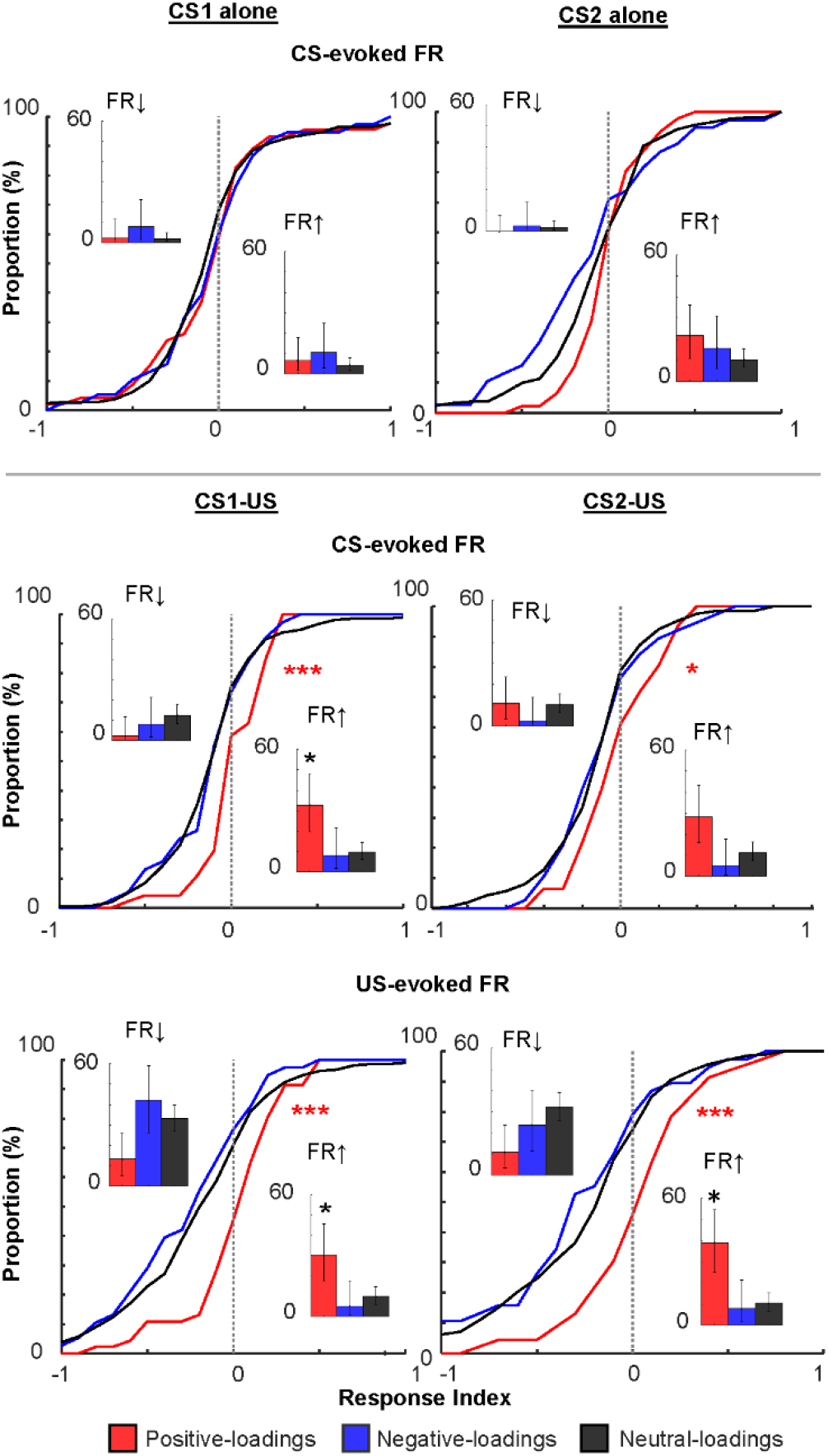
Distributions of the magnitude of stimulus-evoked firing rates. In each neuron, the magnitude of CS- or US-evoked firing rate changes was quantified by calculating Response Index that ranged from −1 to 1. *, *p* < 0.05/2; ***, *p* < 0.001, Kolmogorov-Smirnov test against neutral-loadings neurons. Insets show the proportion of neurons with significant firing rate decrease (left) and increase (right). Error bars show 95% confidence interval. *, *p* < 0.05, Binominal test.

Wilcoxon signed-rank tests were used to test whether a neuron significantly changed firing rates upon CS or US presentations (CS alone blocks, n = 20 trials; CS-US blocks, n = first 50 trials). If the *p*-value was less than 0.05, the neuron was considered as having a significant firing rate increase or decrease in the trial block.

#### Population vector analysis

To examine trial-by-trial changes in ensemble firing patterns (Figures 6A, B), we constructed a population firing rate vector (PV), which contained firing rates of 35 randomly sub-sampled neurons during the CS (from CS onset to US onset; FR_CS) in each trial. The firing rate of each neuron was divided by its maximum firing rate among all trials. PVs were averaged across all trials in each type, and the Pearson correlation coefficient was calculated between these averaged PVs. This procedure was repeated 20 times with different sets of subsampled neurons.

#### Ensemble decoding with support vector machine classifiers

As in our previous study (Morrissey et al., 2017; Xing et al., 2020), we quantified the selectivity of ensemble firing patterns by using decoding accuracy and errors of support vector machine (SVM) classifiers. SVM classifiers produce a model from training data attributes and then predict the target values using only the test data attributes. In our case, the attributes were the normalized binned firing rates of neurons, and the target values were the CS presentation type during which they were recorded. All algorithms were run in Matlab using the freely accessible LIBSVM library that implements the one-against-one approach for multi-class classifications (Chang and Lin, 2011). A firing rate matrix was constructed by concatenating the binned firing rates of a set of 35 neurons during 20 trials randomly drawn, without replacement, from each of four CS presentation types. In each neuron, the firing rate in each trial was divided by the maximum firing rate of the neuron across the 80 trials. The classifiers were trained with the radial basis function (RBF) kernels. Two parameters in the RBF kernel, cost and gamma were first identified by performing a grid search in which five-fold cross-validation accuracy was compared across multiple SVM runs with a different combination of C (11 values) and gamma (10 values). Because the degree of over-fitting should be negatively correlated with the accuracy, we selected the set of C and gamma that resulted in the highest accuracy. These parameters were then used for the training of SVM classifiers with half of the trials (10 trials from each trial type, 40 trials in total). The remaining half was used for testing. The training and testing were repeated 20 times, each of which used a different set of 40 trials for training and another set of 40 trials for testing. Decoding accuracy was defined as the proportion of test trials that were classified correctly. As a measure of generalization of CS1-evoked firing responses to the CS2, we calculated the proportion of errors in discriminating CS2 alone or CS2-US trials from CS1-US trials among incorrectly classified trials.

### Experimental design and statistical analysis

When the measures did not form a normal distribution (Lilliefors test, *p* < 0.05), we used non-parametric tests, including Wilcoxon signed-rank, Wilcoxon rank-sum, Kruskal-Wallis test with *post hoc* Dunn’s tests, or binominal tests. Otherwise, paired t-tests were used. All tests were two-tailed. Sample sizes were determined based upon similar data reported in previous publications. Analyses were performed with Matlab (Mathworks, Natick, MA, United States).

### Histology

Upon completion of all recordings, the location of electrodes was marked by electrolytic lesions. For each tetrode, 5 μA was passed through one wire of each tetrode (positive to the electrode, negative to the animal ground) for 20 s. Rats were then perfused intracardially with 0.9% saline, followed by 10% buffered formalin. The brain was removed from the skull and stored in 10% formalin for several days. For cryosectioning, the tissue was infiltrated with 30% sucrose solution, frozen and sectioned in a cryostat (Leica, Wetzlar, Germany) at 40 μm. Sectioned tissues were stained with cresyl violet and imaged under a light microscope to locate electrode locations. Only recordings from tetrodes located in the prelimbic region were used for the analyses.

## Results

### Novel aversive, but not neutral stimuli induced the abrupt transition of mPFC network state

Seven male rats underwent a sequence of stimulus presentations that were divided into two epochs (Figure 1A). The rats left the conditioning chamber after the first epoch (Epoch 1) and returned the same chamber for the second epoch (Epoch 2). Each epoch included the presentation of one of two neutral stimuli (tone or light, CS1, CS2, 100 ms; Table 1). Initially, the CS was presented by itself every ~30 sec (CS-alone block, ~10 minutes). Subsequently, the CS was paired with a mildly aversive eyelid shock (US, 100 ms) after a short temporal gap (500 ms; CS-US block, ~25-40 minutes). In this paradigm, rats typically require ~seven daily sessions to fully develop anticipatory blinking responses that peak at the expected onset of the US (CR, monitored by eyelid electromyogram; Morrissey et al., 2017; Pilkiw et al., 2017). In the first session, such well-timed CRs were rarely observed; however, a few rats showed some signs of CRs (Figure 1B) and increased the frequency of CR expression (CR%) in the block of CS1-US trials (Figure 1C). As a group, however, the within-session improvement did not reach statistical significance (paired t-test on CR% in 20 CS1 alone trials and the last 20 CS1-US trials, *p* = 0.243). Also, CR% marginally changed across CS2-US trials (*p* = 0.187). When tested next days, these rats significantly increased the frequency of CR expression in the two CS-US blocks (Figure 1C; paired t-test, CS1, *p* = 0.030; CS2, *p* = 0.037), suggesting that the experience during the first day was sufficient to induce lasting changes in behavior. Importantly, this initial sign of memory formation is eliminated by pharmacological inactivation of the prelimbic cortex (Takehara-Nishiuchi et al., 2005). The lack of robust within-session increase in CR expression is likely because the development of specific motor responses (*i.e*., anticipatory blinking responses) takes longer than learning of biological significance of the CS (Rescorla and Solomon, 1967; Lavond et al., 1993; Lee and Kim, 2004).

**TABLE 1.** Types of the stimulus used as the CS in each epoch, Number of recorded neurons, the principal component used for categorizing neurons

| ID | CS1 / CS2 | Number of neurons | Principal component |
| --- | --- | --- | --- |
| Rat 1 | Tone / Light | 30 | PC1 |
| Rat 2 | Light / Tone | 34 | PC1 |
| Rat 3 | Tone / Light | 56 | PC3 |
| Rat 4 | Tone / Light | 41 | PC6 |
| Rat 5 | Tone / Light | 59 | PC6 |
| Rat 6 | Tone / Light | 40 | PC1 |
| Rat 7 | Light / Tone | 35 | PC1 |

During this novel experience, we extracellularly recorded spiking activity from the prelimbic region of the mPFC (Figure 1D). To identify the primary sources of firing rate variance during this novel experience, we applied the principal component analysis (PCA) to binned firing rates (FRs) of simultaneously recorded neurons during the entire recording period, including all trials and inter-trial intervals. In all rats, at least one of the top six principal components (PC) exhibited an abrupt transition upon the first CS1-US trial, which coincided with a drastic firing rate change in a subset of neurons (Figure 2A, PC1 in a representative rat; Table 1). When the rats returned to the same chamber for the second epoch, some of these neurons regained firings and maintained the elevated firing rates throughout the second epoch. In PCA-based 3D projections of ensemble firing patterns (Figure 2B), ensemble activity was clearly separated between the CS1-alone and CS1-US block. The separation of ensemble patterns appeared smaller between the CS2-alone and CS2-US block. To confirm these observations statistically, we calculated the Mahalanobis distances (Hyman et al., 2012; Rozeske et al., 2018) between the binned firing rates of all neurons during the periods before and after the first trial of each type. To eliminate the direct impact of trials on the network activity, bins including stimulus presentations were removed from the analysis. Upon the first CS1-US trial, the ensemble activity changed more robustly than the chance-level change (*i.e*., ensemble differentiation within two adjacent periods in the CS1-alone block; Figure 2C, Wilcoxon signed-rank test; *p* = 0.016). This was not the case after the first CS1-alone trial (*p* = 0.250). Also, in the second epoch, neither type of the first trial induced a significant change in ensemble activity (CS2-alone, *p* = 0.125; CS2-US, *p* = 0.688). Notably, the normalized firing rates averaged across all neurons marginally changed after any of these trials (Figure 2D, Kruskal-Wallis test, all *p*s > 0.05), suggesting that the drastic ensemble transition within a minute after the first CS1-US trial was not due to the sudden changes of firing rate in the entire neural ensembles.

### The network transition was driven by neurons that rapidly developed firing responses selective for stimulus associations

To investigate the single-neuron basis for these population effects, we next examined firing profiles of neurons that had high loadings on the PC with the most robust change after the first CS1-US trial. The loadings of all recorded neurons showed a distribution with heavy tails (Figure 3A; Lilliefors test, *p* < 0.05). Forty-six neurons (15.5%) had loadings greater than or equal to 0.15 (positive-loadings neurons), suggesting that they increased spontaneous firing rates after the first CS1-US trial. In parallel, thirty-eight neurons (12.8%) had loadings less than or equal to −0.15 (negative-loadings neurons), suggesting that they decreased spontaneous firing rates. Spontaneous firing rates were marginally changed in remaining neurons (neutral-loadings neurons).

Over the first few CS1-US trials, positive-loadings neurons as a group increased spontaneous firing rates (*i.e*., firing rates before CS onset) and rapidly developed firing responses to the US (Figure 3B). With the additional few trials, these neurons also developed strong firing responses to the CS1 and reached asymptote in ~9 trials (estimated by fitting an exponential curve; Figure 3C). Once developed, both the CS- and US-evoked firing responses were stable during the remaining CS1-US trials (Figure 3B, C). When the new CS2 was presented, the CS-evoked firing responses were weakened and reached a new asymptote in ~8 trials (Figure 3C, top). When the pairing of CS2 and US began, after a brief period of increase (~2 trials), the response went back to the new asymptote. In contrast, negative-loadings neurons decreased spontaneous firing rates upon the first CS1-US trial and remained silent during the subsequent period (Figure 3B). As evident in the positive value during the CS1-alone block, negative-loadings neurons as a group increased firing rates in response to the CS1 while it was presented by itself (Figure 3C, bottom). The firing responses, however, began to decrease monotonically after the pairing of the CS1 and the US began. They also remained low in the CS2-alone and CS2-US blocks. Neutral-loadings neurons marginally changed baseline and stimulus-evoked firing rates across trials in both epochs (Figure 3B).

Consistent with the visual impression in Figure 3B, positive-loadings neurons showed higher spontaneous firing rates than neutral-loadings neurons in three trial blocks after the first CS1-US trial (Figure 3D, Kruskal-Wallis test, *post hoc* Dunn’s test, all *p*s < 0.001). Importantly, the two neuron types showed comparable spontaneous firing rates before the first CS1-US trial (*i.e*., CS1-alone block; *p* = 0.332). In parallel, the negative-loadings neurons showed a trend toward lower spontaneous firing rates than the neutral-loading neurons starting from the CS1-US block, and the difference reached significance in the CS2-US block (*p* = 0.012).

We also confirmed robust stimulus-evoked firing responses in positive-loadings neurons by calculating the Responsive Index (RI). This measure divided the difference between the stimulus-evoked firing rate and the spontaneous firing rate by the sum of the two. In the two CS-US blocks, the cumulative distribution of the RI of positive-loadings neurons was shifted to the right relative to that of neutral-loadings neurons (Figure 4; Kolmogorov–Smirnov test v.s. neutral-loadings neurons, CS2, *p* = 0.014; other stimuli, *p*s < 0.001). This pattern suggests that positive-loadings neurons were more sensitive to the CS and US than neutral-loadings neurons. Furthermore, compared to the other neurons, the higher proportion of positive-loadings neurons passed a criterion for a significant firing rate increase (insets; binominal test, *p*s < 0.05). In sharp contrast, in the CS-alone blocks, the degree of CS-evoked firing rate changes was comparable between the two neuron types (CS1, *p* = 0.700; CS2, *p* = 0.032), suggesting that positive-loadings neurons increased the sensitivity to the CS only after the CS became predictive of the US. Negative-loading neurons did not show any of these differences from neutral-loading neurons (all *ps* > 0.05).

### Individual neurons developed firing selectivity for stimulus associations within a few minutes

The rapid development of the stimulus-evoked responses was also evident at the level of individual positive-loadings neurons (Figure 5A). Initially, these neurons showed low spontaneous firing rates and did not respond to the CS; however, when the CS-US pairings began, they rapidly developed reliable firing responses to the CS and US within ten pairings (*i.e*., ~5 minutes). In the second epoch, some of these neurons responded to the CS2 immediately (Rat 3, Cell 22; Rat 6, Cell 32) or after it became paired with the US (Rat 1, Cell 10, 30).

**Figure 5.**
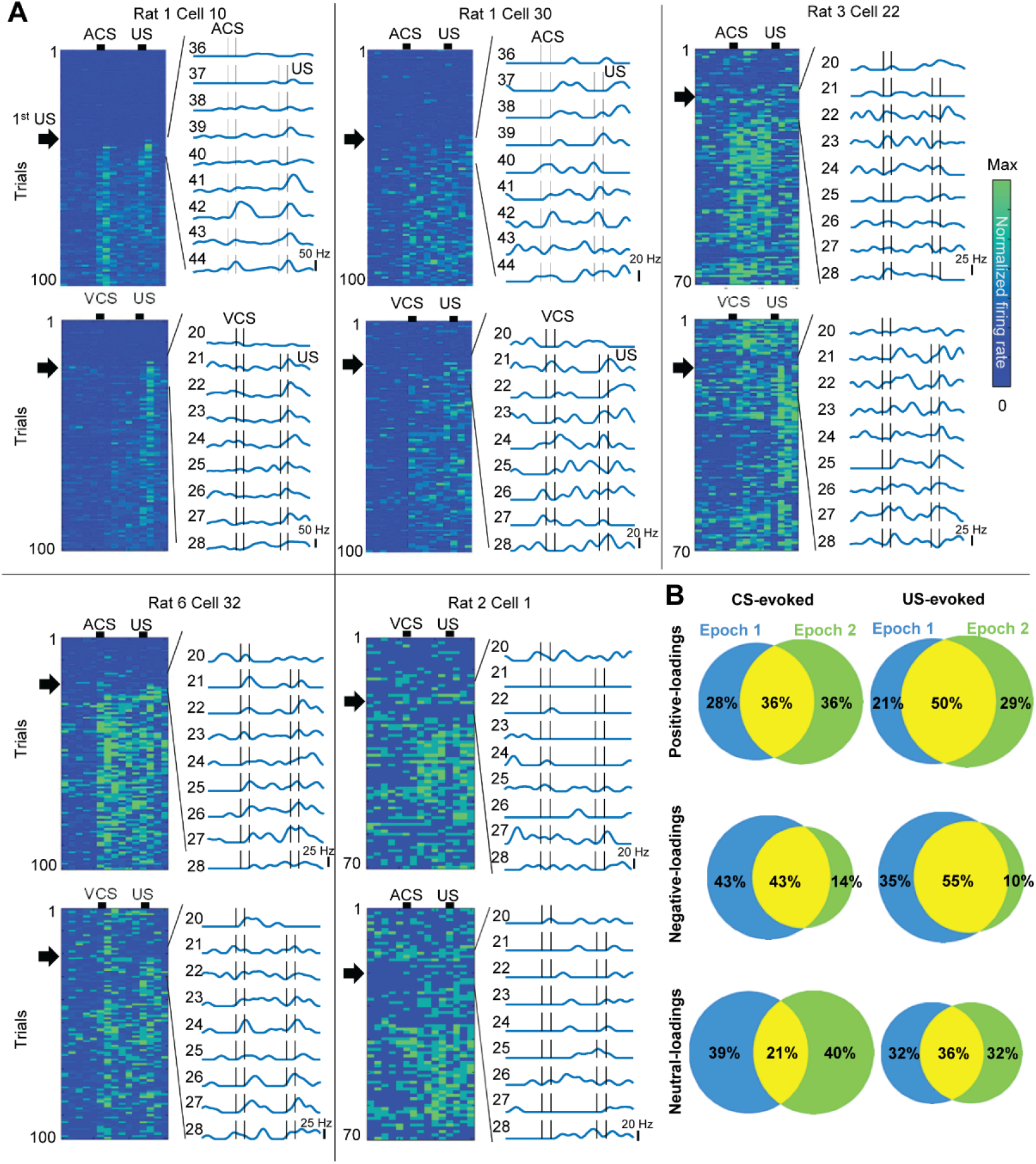
The rapid development of association selectivity in individual neurons. ***A***, Firing rate of individual neurons in all trials (left) and nine trials around the first CS-US trial (Black arrows; right). ***B***, The proportion of neurons with significant firing rate changes upon the CS (left) and US (right) in the first (blue) and second (green) epochs.

Among positive-loadings neurons that significantly changed firing rates in the CS1-US trials, about half of them also changed in the CS2-US trials (Figure 5B). In addition, ~70% of US-responding neurons in the first epoch maintained US-evoked responses in the second epoch. In contrast, the proportion of negative-loadings neurons responding to the CS or US was decreased in the CS2-US block because only a few new neurons acquired firing responses to the CS2 and US.

### Ensemble firing responses were rapidly generalized to different stimuli with the same biological significance

To further examine the degree to which positive-loadings neurons generalized CS1-evoked firing responses to the CS2, we investigated the similarity of CS-evoked firing responses within and between the trial blocks (Figure 6A). When a CS-evoked ensemble firing pattern in one trial (population vector, PV) was correlated against that in another trial of the same type, correlation values were high in all neuron types (within-block consistency, warmer color in white squares). When a CS1-evoked firing pattern was correlated against a CS2-evoked firing pattern, correlation values became low in the CS2-alone block (between-block generalization; cooler color in grey rectangles). Notably, however, only in positive-loadings neurons, the correlation values became high in the CS2-US block (warmer color in a black square). To quantify these observations, we computed correlation coefficients of averaged CS-evoked ensemble firing patterns between two trial blocks (Figure 6B; n = 20 sets of 35 sub-sampled neurons). In the CS2-US block, positive-loadings neurons showed a higher degree of between-block generalization than neutral-loadings neurons (Kruskal-Wallis test, *p* < 0.001; *post hoc* Dunn’s test, *p* = 0.002). This was not the case in the CS2-alone block (*p* = 0.056). Furthermore, the within-block consistency was higher in positive-than neutral-loadings neurons (*p* < 0.001). In parallel, negative-loadings neurons showed comparable within-block firing consistency (*p* = 0.930) to neutral-loading neurons. Consistent with gradual loss of CS-evoked firing responses (Figures 3B, 3C, 5B); however, they showed lower between-block firing generalization to the CS2 (CS2-alone, *p* < 0.001; CS2-US, *p* = 0.001).

**Figure 6.**
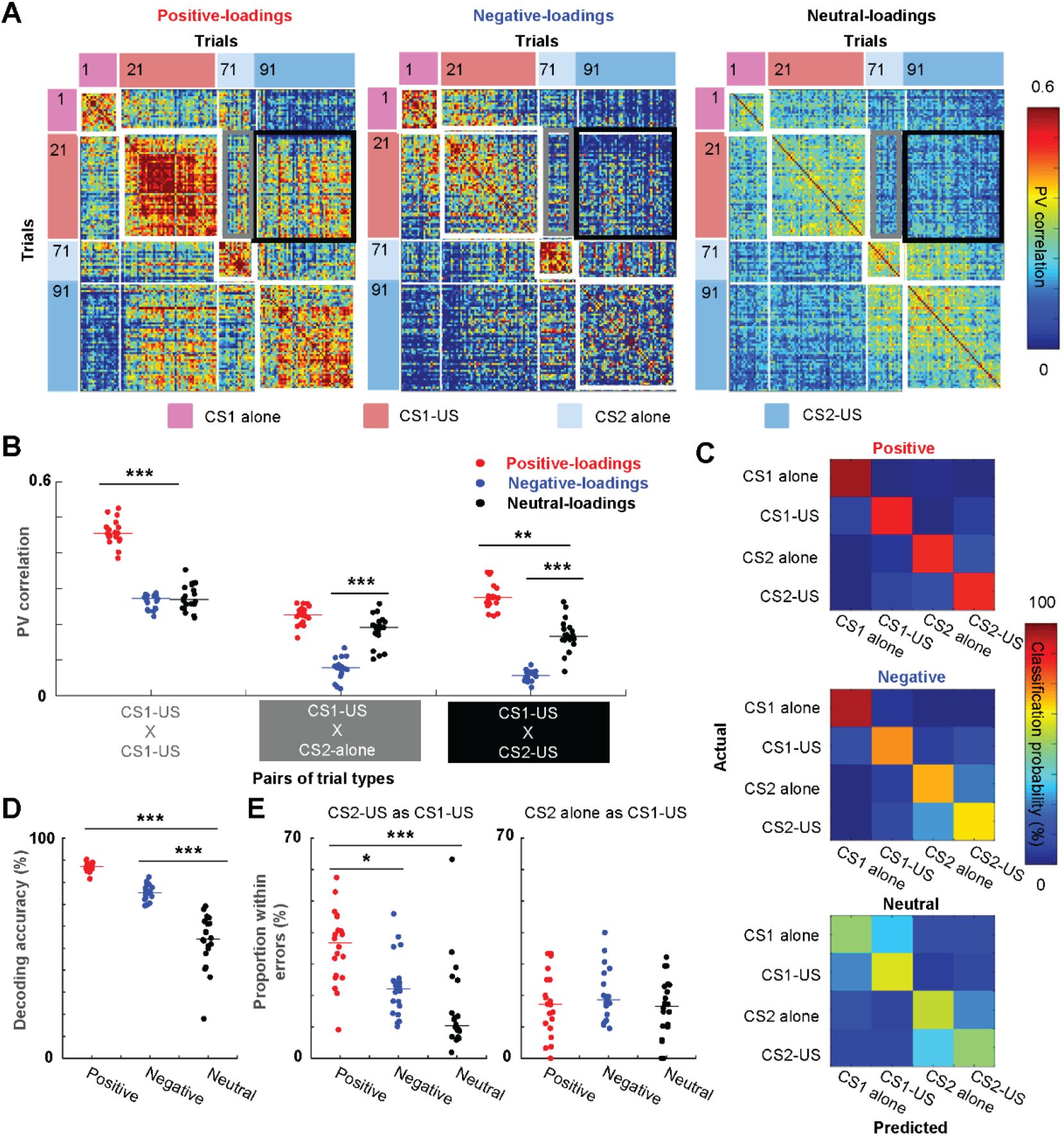
Generalization of ensemble firing patterns to a new stimulus with the same biological significance. ***A***, Pearson correlation coefficients of CS-evoked ensemble firing vectors (PV) between all possible pairs of trials. ***B***, The correlation values (n = 20 sets of 35 sub-sampled neurons; lines show median) between the averaged PV in the CS1-US block and that in the same (within-block consistency) or other blocks (between-block generalization). ***C***, Representative confusion matrices from support vector machine (SVM) classifiers showing the probability (color) that a trial of one type (rows) was classified as another type (columns). ***D***, Decoding accuracy of SVM classifiers applied to 20 sets of 35 sub-sampled cells (lines show median). Decoding accuracy was significantly higher with positive-loading neurons than neutral-loading neurons. ***E***, The proportion of the specific type of classification errors in a total number of misclassified trials (n = 20 sets of 35 sub-sampled neurons; lines show median). **, *p* < 0.01; ***, *p* < 0.001, Kruskal-Wallis test, *post hoc* Dunn’s test.

As another measure for ensemble selectivity, we also used support vector machine (SVM) classifiers to quantify the degree to which ensemble firing patterns differentiated trials of each block (Morrissey et al., 2017; Xing et al., 2020). Better performance of the classifier reflects higher selectivity of CS-evoked firing patterns for the modality of the CS (tone or light) and its association with the US (presented alone or paired with the US). In parallel, the probability of errors in discriminating CS1 and CS2 trials indicates the degree to which neuron ensembles generalized CS1-evoked firing patterns to the CS2. Examination of the classifier outputs (the ‘confusion matrix,’ Figure 6C) showed that positive-loadings neurons successfully differentiated all four CS types, while neutral- and negative-loadings neurons differentiated CS1 and CS2 trials better than CS-alone and CS-US trials. Compared to neutral-loadings neurons, the overall decoding accuracy was significantly higher in positive- and negative-loadings neurons (Figure 6D; n = 20 runs with 35 sub-sampled neurons; Kruskal-Wallis test, *p* < 0.001; *post hoc* Dunn’s test, *ps* < 0.001). When the input was CS2-US trials, some inaccurate classifications resulted from errors in discriminating them from CS1-US trials, and the proportion of this error type was significantly higher in positive-loadings neurons than the other neurons (Figure 6E, left; 100% = total errors; Kruskal-Wallis test, *p* < 0.001; *post hoc* Dunn’s test, v.s. neutral, *p* < 0.001; v.s. negative, *p* = 0.024). In contrast, when the input was CS2-alone trials, the proportion of errors in discriminating them from CS1-US trials were comparable across three neuron types (Figure 6E, right; Kruskal-Wallis test, *p* = 0.470). These results suggest that positive-loadings neurons formed highly stable firing responses to the CS1 and generalized the responses to the CS2 only after it became associated with the same US as the CS1.

## Discussion

Theories posit that the hippocampus rapidly forms associations among ongoing events as they unfold and later instructs the gradual stabilization of their memory traces in the neocortex.

Consistent with this two-stage model of memory consolidation, over weeks after learning, the mPFC undergoes various genetic, physiological, and structural changes necessary for stabilizing memory traces. Parallel evidence, however, suggests that these changes might be initiated at the time of encoding, leading to a view that the coordinated formation of memory traces within hippocampal-neocortical circuits at the time of encoding. In support of the latter view, we found that ~15% of prelimbic neurons underwent two types of functional changes during a novel experience, 1) abrupt increase in spontaneous firing rates within a minute after the first shock presentation and 2) subsequent development of firing responses selective for the stimulus-shock association within ~ten instances of their pairings.

The group of prefrontal neurons showed an abrupt, coordinated change in firing rates within a minute after a single aversive stimulus that converted a seemingly neutral experience to behaviorally relevant experience (Figures 2A-C). Such rapid changes did not occur upon a novel, neutral stimulus that was presented by itself, suggesting that the negative valence, but not the novelty of stimulus triggered the network transition. Furthermore, once developed, the new network state was maintained stably during the rest of the experience, making the present observation distinct from gradual firing rate changes tracking elapsed time in the mPFC (Hyman et al., 2012) or other brain regions (Mankin et al., 2015; Tsao et al., 2018; Diehl et al., 2019). The sudden shift in the network state bears a resemblance to prefrontal ensemble firing patterns when *well-trained* rodents experienced a switch between previously learned behavioral contexts, such as sets of action-outcome contingency (Karlsson et al., 2012), sequences of action (Ma et al., 2014), and the combination of environmental features signaling threat (Rozeske et al., 2018). Our data extend these observations by uncovering that the sensitivity to a relevant behavioral context is an innate property of the mPFC network.

The observed, abrupt network transition was driven by ~15% of neurons that immediately elevated spontaneous firing rates upon the first US presentation (Figures 3A, B, D). Moreover, these, but not the other, neurons rapidly developed firing responses to the preceding neutral stimulus (CS) with a few instances of CS-US pairings (Figures 3B, C, 4, 5A). Once developed, CS-evoked firing rates were stable across repeated stimulus pairings (Figures 3B, C, 5A), which sharply contrasts with a gradual decrease in CS-evoked firing patterns of neurons in the caudal anterior cingulate cortex (Weible et al., 2003; Bryden et al., 2011; Hattori et al., 2014) associated with attention to cues. Similarly, the US-evoked firing rates were also stable across repeated CS-US paired trials (Figures 3B, 5A), which is different from the prediction error signal for aversive outcome reported in the periaqueductal grey (Johansen et al., 2010) and sub-regions in the ventral tegmental area (Menegas et al., 2018).

Notably, the neurons that rapidly acquired firing responses to the CS-US association were indistinguishable from the remaining neurons in terms of the initial responsiveness to the CS before it was paired with the US (Figure 4). This observation suggests that at baseline, they did not receive a greater amount of CS inputs than the remaining neurons and contradicts the traditional Hebbian plasticity model of associative learning: among cells receiving the CS signals, those activated by the US will potentiate their CS-evoked responses (Blair et al., 2001; Medina et al., 2002; Maren and Quirk, 2004). Alternatively, the emerging idea is that neural responsiveness can also be amplified by the plasticity of intrinsic neuronal excitability (Titley et al., 2017; Lisman et al., 2018; Debanne et al., 2019). The elevated intrinsic excitability enables small synaptic inputs, previously ineffective, to gain the ability to drive the initiation of action potentials (Haider and McCormick, 2009; Stuart and Spruston, 2015). Although we did not measure intrinsic excitability directly, the abrupt, sustained increase in spontaneous firing rates within a minute after the first CS-US pairing (Figures 3B, D) supports this view. Future investigations with *in vivo* intracellular recording are necessary to probe the link between intrinsic plasticity and memory-selective neural firings in the mPFC.

Among positive-loadings neurons that acquired firing responses to the CS1, ~50% of them also acquired firing responses to the CS2 (Figure 5B). In line with this observation, two independent quantifications of ensemble selectivity revealed that positive-loadings neurons generalized CS1-evoked ensemble patterns to the CS2 (Figures 6B, E). One may argue that the rapid firing generalization was simply due to the elevated sensitivity to any sensory stimuli. This, however, is unlikely because the generalization of firing responses became prominent after the CS2 became paired with the US (Figures 6B, E). In fact, the similarity of firing responses between CS2-alone and CS1-US trials was comparable among three neuron types (Figures 6B, E). The present observations, therefore, suggest that the mPFC network is capable of discriminating the CS2 from the CS1 but actively assimilates the ensemble codes for the two stimuli after they became associated with the same biological significance.

We previously reported similar firing assimilation occurring over weeks after learning and argued that this unique circuit property enables the mPFC to extract commonality from multiple similar experiences (Morrissey et al., 2017). Theories posit that these pattern recognition and integration processes allow for building an internal model of the world, in other words, prior knowledge or schema (Marr, 1971; McClelland et al., 1995; Winocur et al., 2010; Sekeres et al., 2018). The present finding raises the possibility that the mPFC might also play a similar role at the time of memory encoding (Takehara-Nishiuchi, 2020b). Rapid firing generation from the CS1 to the CS2 alludes to the integration of stimulus information with the same biological significance over time, which enables the mPFC to detect, in real time, relevant contents of a new experience. In doing so, it may emit a relevancy signal that enhances the contrast between central and incidental contents in hippocampal memory traces. The close mPFC-hippocampal interaction during encoding also ensures that newly formed traces in the mPFC and hippocampus are liked with one another, thereby facilitating the subsequent, progressive rewiring of memory networks during systems consolidation.

In summary, the present findings identified experience-dependent neural plasticity in the mPFC that develops at a comparable speed to that reported in the hippocampus (Whitlock et al., 2006; Bittner et al., 2015, 2017; Broussard et al., 2016) and amygdala (Rosenkranz and Grace, 2002; Herry et al., 2008; Grewe et al., 2017; Zhang and Li, 2018). The rapid plasticity enables a subset of the mPFC neurons to develop firing responses selective for behaviorally relevant content during experiences, thereby providing evidence for fast, perhaps more intelligent learning taking place in the neocortex. Future studies must investigate whether the rapid neocortical plasticity depends on the integrity of the hippocampus and how it relates to online, neocortical-hippocampal interaction during encoding (Shin et al., 2019).

## ACKNOWLEDGMENTS

This work was supported by NSERC Discovery Grant, CFI Leaders Opportunity Fund (K. T.-N.), and NSERC graduate fellowship (M.M., M.P.). The authors thank Andrew Peluso for technical assistance.

